# Targeting B and T lymphocyte attenuator regulates lupus disease development in NZB/W mice

**DOI:** 10.1101/2024.05.28.596218

**Authors:** Léa Gherardi, Lucie Aubergeon, Mélanie Sayah, Jean-Daniel Fauny, Hélène Dumortier, Fanny Monneaux

## Abstract

B and T Lymphocyte Attenuator (BTLA) is a co-inhibitory receptor expressed by most immune cells, playing a role in negatively regulating immune responses. Studies in MRL/lpr lupus mice deficient for BTLA, indicate that BTLA has a protective role in lupus. We have previously shown an altered BTLA expression by regulatory T cells and an impaired capacity of BTLA to inhibit CD4^+^ T cell activation in lupus patients. In this study, we thoroughly analyzed BTLA expression and function in the NZB/W lupus-mouse model. We found that diseased NZB/W mice exhibit a BTLA expression and function pattern similar to that observed in lupus patients, emphasizing the importance of this mouse model in evaluating the therapeutic potential of targeting BTLA. Administration of a monoclonal anti-BTLA antibody (clone 6F7, which displays agonist properties *ex vivo*) into pre-diseased NZB/W mice resulted in a delayed onset of proteinuria, limited kidney damages and an increased survival rate compared to isotype-treated mice. This beneficial effect was associated with a decrease in circulating B cell frequency and required continuous exposure to the antibody. Regarding its mode of action, we demonstrated that the 6F7 antibody is not a depleting antibody and does not block HVEM binding to BTLA, but instead induces BTLA down modulation, leading to a selective reduction of follicular B cell numbers, and exhibits *in vivo* agonist activity. Overall, our data confirm the involvement of BTLA in lupus pathogenesis and provide the first evidence that BTLA is a potential therapeutic target for the treatment of lupus.

## INTRODUCTION

B and T lymphocyte attenuator (BTLA) is an inhibitory receptor of the CD28 superfamily that negatively regulates the immune response in synergy with CTLA-4/B7 and PD-1/PDL1 pathways (1). BTLA is expressed on T cells (CD4^+^ and CD8^+^ T cells), B cells, but also natural killer (NK) cells, NKT cells, macrophages and dendritic cells (DC) (2). Moreover, BTLA was described to be expressed by T follicular helper (T_FH_) cells (3). The ligand for BTLA is Herpes Virus-Entry Mediator (HVEM; TNFRS14) (4), a TNF receptor family protein found on T, B, NK, DC and myeloid cells. The ligation of BTLA by HVEM attenuates T-cell activation, leading to decreased cell proliferation, cytokine production and cell cycle progression. The inhibitory role of BTLA *in vivo* was revealed by the analysis of BTLA-deficient mice characterized by a breakdown of self-tolerance, resulting in the development of an autoimmune hepatitis-like disease, lymphocytic infiltration in multiple organs (5) and enhanced specific antibody responses and sensitivity to EAE (1). On the contrary, BTLA engagement through the use of agonist antibodies leads to the induction of tolerance. Indeed, the administration of an agonist anti-BTLA antibody 3C10 was shown to prolong heart allograft survival via the induction of IL-10-producing regulatory T cells (6) and to prevent atherosclerosis by shifting the balance between follicular B cells and regulatory B cells (7). Moreover, the agonistic BYK1 anti-BTLA antibody prevents the development of graft versus host disease (GVHD), if it was administered at the time of transplantation (8). BTLA is thus considered as a critical receptor for maintaining peripheral tolerance.

Systemic lupus erythematosus (SLE) is a highly complex and heterogeneous autoimmune disease characterized by the presence of autoantibodies directed against a spectrum of self-antigens (including nuclear antigens) and by the subsequent formation of immune complexes (9).

Hallmark features of SLE are immune system activation and inflammation in multiple organs, such as the skin, joints, kidneys, lungs and central nervous system. As autoantibody producers, B cells are attractive targets to treat lupus (10). However, clinical trials aiming at depleting B cells, revealed disappointing results, probably because plasma cells do not express typical B cell surface markers such as CD20 or CD22. Thus, modulation of plasma cell differentiation by targeting T-B dialog may be an interesting alternative for lupus patients who are refractory to B cell depletion (11).

Interestingly, the lupus disease is exacerbated in MRL/lpr lupus mice deficient for BTLA (12) suggesting a protective role of BTLA in lupus. We previously investigated the expression of BTLA on T cell subsets in lupus patients and described that the enhancement of BTLA expression after activation is significantly lower in SLE patients compared to healthy donors (HD). In addition, we showed that there is a significant decrease in the capacity of BTLA to inhibit the proliferation and the activation of lupus CD4^+^ T cells compared to HD. We evidenced that BTLA deficiency in lupus settings is due to the poor BTLA recruitment to the immunological synapse following activation, and that this can be corrected by restoring intracellular trafficking (13). Moreover, we very recently observed that BTLA expression is significantly increased on activated Tregs (aTregs) from lupus patients, especially those displaying an active disease (14). BTLA expression by lupus aTregs correlates with the diminution of the aTregs frequency observed in these patients, suggesting that the higher BTLA expression on the surface of lupus aTregs may account for the reduced frequency of this Treg subset in lupus patients. Altogether, our results highlight a possible involvement of defective BTLA signaling pathway in lupus patients. As mouse models are crucial for designing and evaluating therapeutic protocols, we now wish to investigate BTLA expression and function in lupus mice.

In the present study, we characterized the phenotype and function of BTLA expressing cells in NZB/W F1 mice, that spontaneously develop a syndrome resembling human SLE notably because of the development of severe immune complex-related glomerulonephritis (15). We also assessed the *in vivo* effects of an anti–BTLA antibody administration in lupus NZB/W F1 mice. We found that diseased-NZB/W mice display similar pattern of BTLA expression and function than the one we previously observed in human SLE patients, highlighting the relevance of this mouse-model for evaluating the therapeutic potential of BTLA targeting. Very interestingly, we demonstrated that an anti–BTLA antibody was effective for alleviating the lupus-like nephritis of NZB/W F1 mice.

## MATERIALS AND METHODS

### Mice

Female BALB/W (BALB/c x NZW) F1 and NZB/W (NZB x NZW) F1 mice were bred in our animal facilities (approved by French Veterinary Services, #I-67-482-2). All experiments were carried out in conformity with the 2010/63/UE European animal bioethics legislation and were approved by the Regional Ethics Committee of Strasbourg (CREMAS) and by the French Ministry of Higher Education and Research (APAFIS #2020041717124974).

### Treatment of mice

NZB/W females (20-22 weeks of age) were randomly assigned to treatment groups and received either an i.p. administration of the anti-BTLA antibody (clone 6F7; 3mg/kg/injection, BioXcell, Lebanon, NH, USA), or an appropriate mouse isotype antibody (IgG1κ; 3mg/kg/injection, BioXcell,) twice a week for 10 weeks. Urine protein levels and weight were determined twice a week and mice exhibiting high proteinuria (score >3) for at least two consecutive weeks were considered to be severely proteinuric. Proteinuria was evaluated in fresh urine samples using a colorimetric assay for albumin (Albustix, Siemens, Munich, Germany) and was semi-quantitatively estimated according to the scale recommended by the manufacturer. Blood samples were collected every 3 weeks to evaluate auto-antibody levels in the serum and to perform a phenotypic analysis in peripheral blood. Mice were euthanized either *i*) at limit point achievements (high proteinuria and/or 20% weight loss and/or prostration) in order to evaluate clinical and biological signs of the disease (proteinuria, autoantibodies, survival) or *ii*) at 33 weeks of age to evaluate the therapeutic effect of anti-BTLA administration on kidney damages.

### Depletion of NK cells and macrophages

NZB/W were treated with an anti-NK1.1 antibody (clone PK136, Biolegend, Amsterdam, The Netherlands; 200 µg i.p. on days −2, 0 and 3) to deplete NK cells. Mice were then administered with a high dose of the anti-BTLA 6F7 antibody (500 µg/mouse, i.p.) on day 0, and were sacrificed on day 5. For macrophages depletion, mice were treated with clodronate-loaded liposomes (Liposoma BV, Amsterdam, The Netherlands; 200 µl/mouse, i.v.) one day before the 6F7 administration.

### Cell isolation and cell culture

Total or naive CD4^+^ T cells were negatively isolated from spleen using Mojosort Mouse CD4 T Cell Isolation Kit or Mojosort Mouse CD4 Naive T Cell Isolation Kit (Biolegend) according to the manufacturer’s instructions. The purity of the total CD4^+^ or naive CD4^+^ T cell population was typically ≥ 90%. Cells were cultured in complete medium (RPMI 1640 containing 10% FCS, 10 µg/ml gentamicin, 0.05 mM ß-mercaptoethanol and 10 mM HEPES) at the rate of 100 000 cells/well in 96-well plates at 37°C.

For leucocytes isolation from kidneys, kidneys were cut and digest for 30 min at 37°C with 0.5mg/ml collagenase D (Roche, Mannheim, Germany) and 0.05 mg/ml DNase I (Roche) in complete medium. Tissue homogenates were sequentially filtered through 100-, 70- and 40-µm nylon meshes and washed with complete medium. Leucocytes were isolated using a 72/36% discontinuous Percoll gradient (Sigma-Aldrich, Saint Louis, USA) and centrifugation (30 min, 400g). The leucocyte-enriched cell suspension was harvested from the Percoll interface.

### BTLA functional analysis in CD4^+^ T cells

Purified CD4^+^ T cells or naive CD4^+^ T cells were stimulated with beads (Dynabeads M-450 Epoxy, ThermoFischer Scientific, Waltham, USA) coated with anti-CD3 (7.5%, clone 145-2C11, BD Biosciences, San Diego, USA) / anti-CD28 (7.5%, clone 37.51, BD Biosciences) / anti-BTLA (85%, clone 6F7, ThermoFischer Scientific) or IgG1κ (85%, clone P3.2.8.2) antibodies (ratio bead/cell = 1:1). Flow cytometric analysis of CD69, CD25 and BTLA was performed after 24 hours and 48 hours of culture. To measure cell proliferation, cells were incubated for 10 min at 37°C with 5 µM CellTrace Violet (ThermoFisher Scientific). After washing, cells were activated as describe above and cultured for 5 days.

### BTLA functional analysis in B cells

Splenocytes from mice that had received either the anti-BTLA 6F7 antibody or the isotype control (3mg/kg/mouse, i.p. on day 0 and day 4) were collected on day 6 and stimulated with a polyclonal anti-IgM antibody (10 µg/mL, Jackson, West Grove, USA) and a polyclonal anti-IgG antibody (20 µg/mL, Jackson). Flow cytometric analysis of CD69 was performed after 6 hours of culture.

### Flow cytometry analysis

Cells isolated from various tissues (spleen, blood, kidneys) were stained for 20 minutes at 4°C in staining buffer (2% FCS in PBS) with following conjugated antibodies: CD4 (clone RM4-5), CD8 (clone 53-6.7), CD19 (clone 1D3), CD23 (clone B3B4), CD25 (clone PC61), CD62L (clone MEL-14), CD69 (clone H1.2F3), CD138 (clone 281-2) from BD Biosciences, CD3 (145-2C11), CD44 (clone IM7), CD45 (clone 30F11), B220 (clone RA3-6B2), BTLA (clone 8F4) from Biolegend, BTLA (clone 6F7), CD21 (clone 8D9), Foxp3 (clone FJK-16s) from ThermoFischer Scientific. For mouse blood phenotyping, 50 µl of blood were first incubated with conjugated-antibodies for 20 minutes at 4°C in staining buffer (2% FCS in PBS). Red blood cells were then lysed with lysis buffer (EasyLyse, Agilent Technologies, Santa Clara, USA) and cells were washed twice in staining buffer. Absolute cell numbers were determined in the blood using Precision Count Beads (Biolegend). Apoptosis was evaluated thanks to 4’,6-diamidino-2-phenylindole (DAPI; Life Technologies, Carlsbad, USA) and Annexin-V-APC (BD Biosciences) staining. HVEM binding to BTLA was assessed by incubating cells with the HVEM-Fc recombinant protein (20 µg/ml; Biolegend) followed by a staining with an Alexa488 conjugated-anti-IgG1 secondary Ab (clone HP6069, ThermoFischer Scientific). Cells were first incubated with rat anti-mouse CD16/32 monoclonal Ab (2.4G2, BioXcell) to block Fcγ receptors. For FoxP3 and BTLA intracellular staining, cells were fixed and successively permeabilized (FoxP3/staining buffer eBiosciences™ or BD Cytofix/Cytoperm™ and BD Cytoperm Permabilization Buffer Plus™) according to recommended instructions. Dead cells were excluded using DAPI and single cells were discriminated from aggregates or doublets using SS-W *versus* SS-H and FS-W *versus* FS-H plots. Cell acquisition was performed using 10-color Flow Cytometer Gallios-Navios (Beckman Coulter, Brea, USA). Data were analyzed using FlowJo 10.4 software (Treestar, Oregon, USA).

### Assessment of kidney disease

For histology, kidneys were fixed overnight at 4°C in paraformaldehyde 4% and embedded in paraffin (Leica, Wetzlar, Germany). Sections of 5 µm were dewaxed, rehydrated and stained with hematoxylin and eosin (H&E), dehydrated and permanently mounted. Pathological changes in the kidneys were assessed by evaluating glomerular and tubular activity (i.e., increased glomerular size, hyalin deposit, glomerular lesions and tubulointerstitial infiltration). Sections were scored using a 0-4 scale as follows: 0 = no lesion, 1 = lesions in <25% of glomeruli/tubules, 2 = lesions in 25-50% of glomeruli/tubules, 3 = lesions in 51-75% of glomeruli/tubules and 4 = lesions in > 75% of glomeruli/tubules. Images were acquired with a 40x objective on a NanoZoomer S60 (Hamamatsu Photonics, Japan) and analyzed with the QuPath Software (16).

For immunofluorescence, kidneys were frozen in Tissue-Tek® O.C.T™ Compound ( CellPath, Mochdre, United Kingdom) and 7µm-sections were prepared. They were then mounted on SuperFrost™ Plus glass slides (ThermoFischer Scientific) fixed in cold acetone for 20 min, air dried, and washed in PBS. Sections were saturated with BSA 2% PBS and then incubated overnight with an APC-conjugated anti-CD45 antibody (clone 30F11, Biolegend) at 4°C. After fixation with paraformaldehyde 4% (Sigma-Aldrich) for 20 min, slides were then stained with DAPI (100 ng/ml) for 15 min at room temperature and mounted with Invitrogen Fluoromount-G Mounting Medium (ThermoFischer Scientific). Mosaic images were acquired with a 20x objective on a Zeiss Axiovert Z1 driven by Metamorph software (Molecular devices). Stitching of images was performed on Metamorph Software and quantification of infiltrate area was performed using both ImageJ (17) and Ilastik (18) softwares.

### Statistical analyses

Data are shown as mean +/-SEM. Data were analyzed using a Mann-Whitney or Kruskal-Wallis test. Data concerning survival or proteinuria were analyzed using a Log-Rank test (Prism version 6.0; GraphPad Software). P-values <0.05 were considered significant.

## RESULTS

### BTLA expression in lupus NZB/W mice

To determine the role of BTLA in lupus pathogenesis in the NZB/W mouse model, we first examined BTLA expression on B and T lymphocyte subsets at various stages of the disease, i.e. prior biological and clinical symptoms appearance (10-12 week-old) and after disease development (mice older than 35 weeks and displaying proteinuria), compared to haplotype-matched BALB/W ([BALB/c X NZW] F1 carrying the H2^d/z^ haplotype and the same BTLA.1 allele as NZB/W mice). As described in other mouse strains, we observed that BTLA expression is higher on B cells than on both CD4^+^ and CD8^+^ T cells of young BALB/W mice (Fig. 1A, p<0.0001). Naive (CD44^-^CD62L^+^) and effector/memory (CD44^+^CD62L^-^) CD4^+^ T cells express similar levels of BTLA whereas in B cells, BTLA expression is significantly higher on follicular B cells (FO; CD21^+^CD23^+^) compared to marginal zone B cells (MZ; CD21^+^CD23^-^) and newly formed CD21^-^CD23^-^ B cells (Fig. 1A). When we compared BTLA expression levels between BALB/W and NZB/W mice, we found that BTLA is not differentially expressed on CD4^+^ T cells (or CD4^+^ T cell subsets) nor CD8^+^ T cells compared to control age-related BALB/W mice (Fig. 1B, C). Concerning B cells, BTLA is similarly expressed by total CD19^+^ B cells from NZB/W mice and BALB/W mice (Fig. 1B), however, we observed a significant increase of BTLA on CD21^-^CD23^-^ B cells from lupus mice whatever the age (p<0.01 for young mice, p<0.05 for old mice), and a higher BTLA expression on MZ B cells from old-diseased NZB/W mice compared to old BALB/W mice (p<0.01; Fig. 1D). Finally, we confirmed that BTLA is expressed by NZB/W plasma cells (Fig. 1E). Altogether, our results show that BTLA expression is only slightly modified in NZB/W lupus mice compared to age related BALB/W.

**Figure 1:**
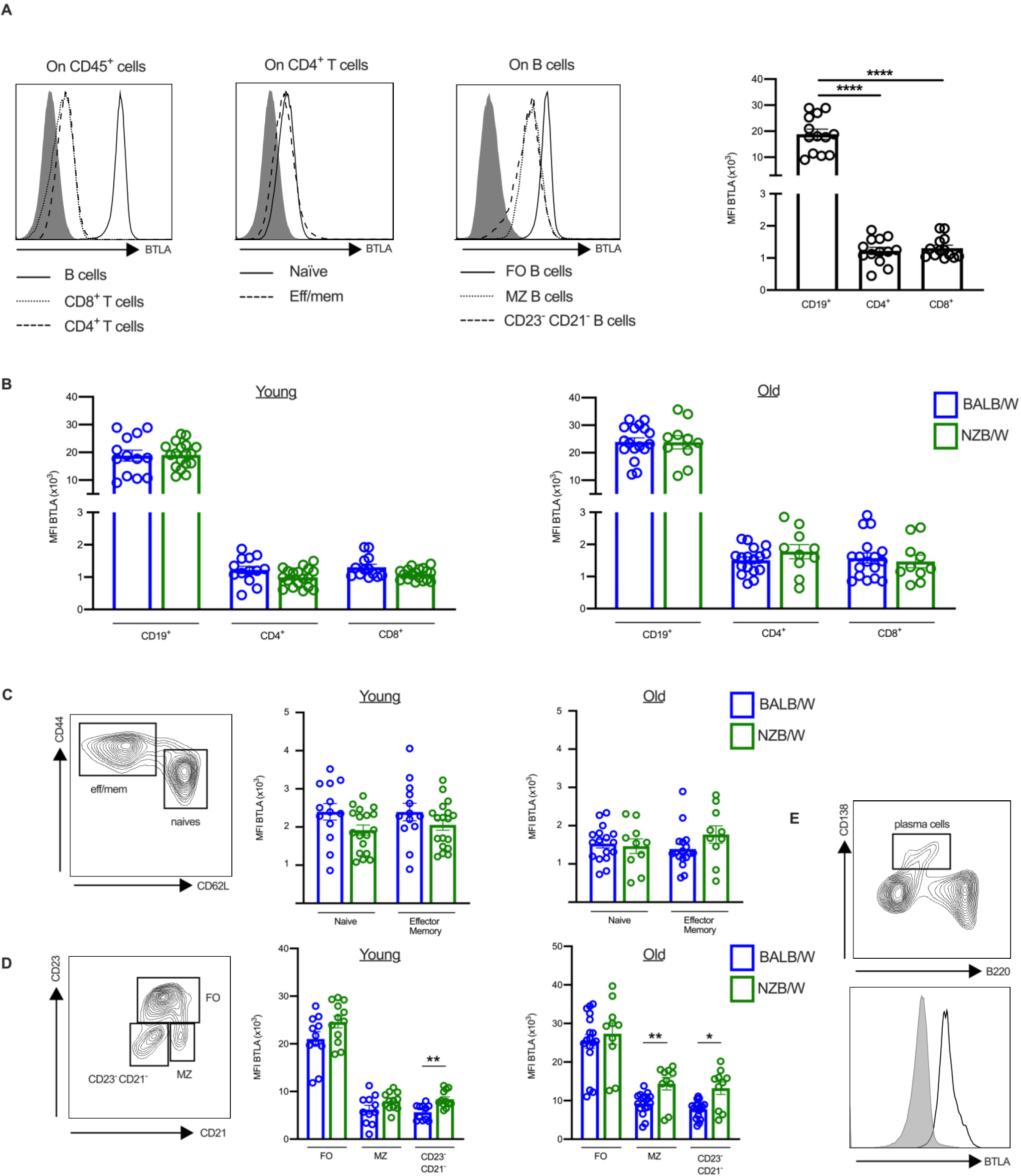
BTLA expression on T and B cell subsets. **(A)** Flow cytometry analysis of BTLA expression on spleen lymphocytes from young BALB/W mice (n=13). **(B-D)** Comparison of BTLA expression on CD19^+^ B cells, CD4^+^ and CD8^+^ T cells (**B**), CD4^+^ T cell subsets (**C**), and B cell subsets (**D**) in young or old BALB/W (blue bars, n=13 and n=17), and young or old NZB/W (green bars, n=18 and n=10). **(E)** BTLA expression on plasma cells in old NZB/W mice. Results are expressed as mean ± SEM and each dot represents one mouse. *p<0.05; **p<0.01; ****p<0.0001, Welch’s ANOVA (A), Mann-Whitney test (B-D).

### BTLA function in lupus NZB/W mice

As we previously evidenced a defective BTLA signaling pathway in human lupus CD4^+^ T cells, we wondered whether the engagement of BTLA could efficiently inhibit CD4^+^ T cell activation in lupus mice. The only studies in which anti-BTLA antibodies were used in “agonist tests” *in vitro* were performed in the C57BL/6 mouse model bearing the *Btla^b^* encoding BTLA.2 variant, whereas NZB/W mice express the BTLA.1 variant (encoded by *Btla^a^*) (19). We first confirmed that the anti-BTLA 6F7 clone (specific for both BTLA.1 and BTLA.2 alleles; agonist activity in the C57BL/6 mouse model), also possesses agonist properties in mice carrying the BTLA.1 allele. Indeed, CD4^+^ T cell activation is inhibited in BALB/W when TCR and BTLA are co-engaged as shown by the expression of the two activation markers CD69 and CD25 (Supplementary Fig. 1). In contrast, we observed a decreased capacity of BTLA to inhibit CD69 (p<0.001) and CD25 (p<0.0001) expression in old-diseased NZB/W mice compared to BALB/W mice of the same age (Fig. 2A). Accordingly, the inhibition of the proliferation is also defective in old-diseased lupus mice (p<0.01) (Fig. 2A).

**Figure 2:**
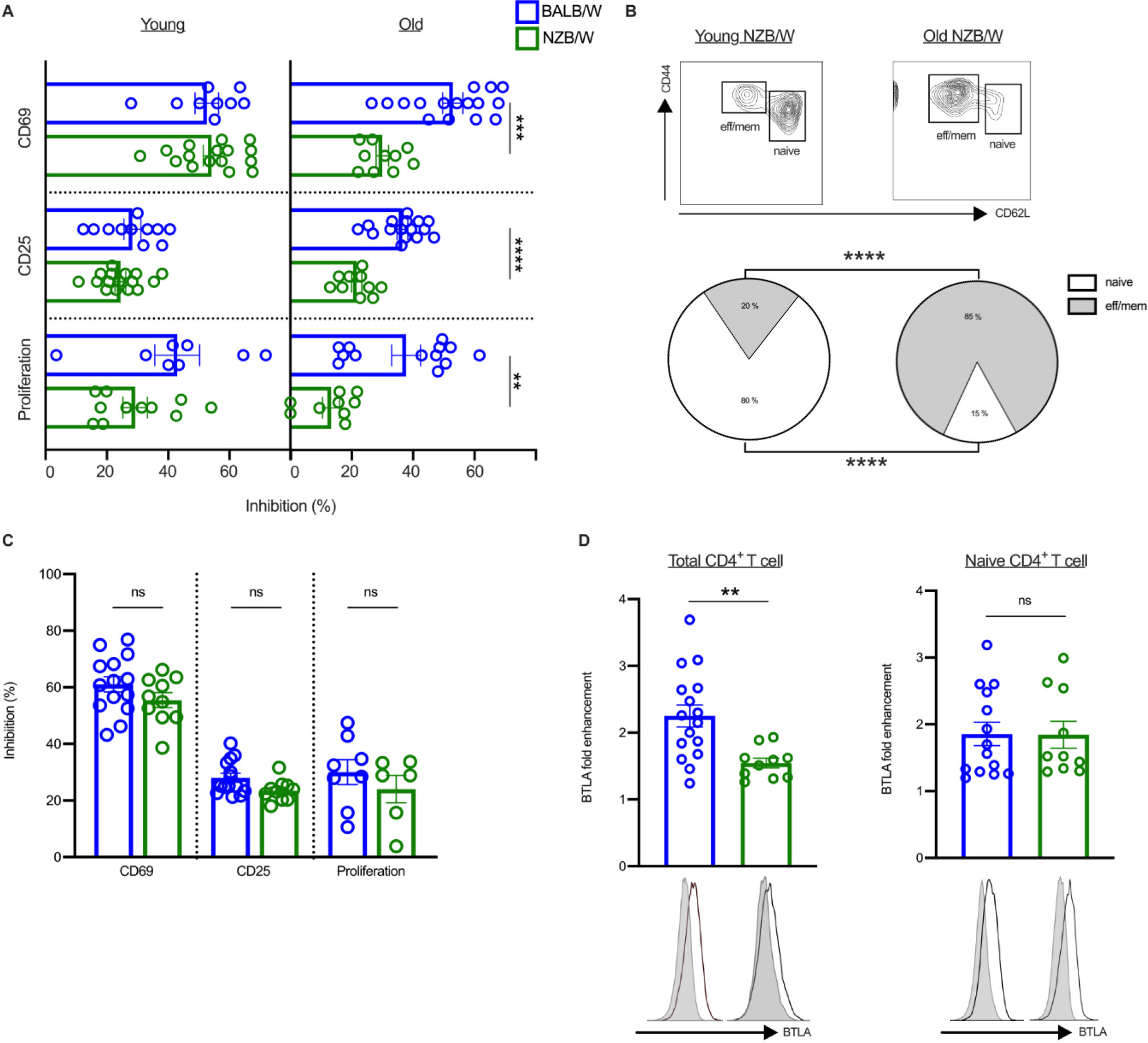
Impaired BTLA functionality in CD4^+^ T cells from old NZB/W mice. **(A)** Purified CD4^+^ T cells from young or old BALB/W (blue bars, n=8-16) and young or old NZB/W (green bars, n=9-16) were stimulated with anti-CD3/anti-CD28 antibody-coated beads in the presence of the agonist anti-BTLA 6F7 antibody or its isotype control. The expression of CD69 at 24h and CD25 at 48h were analyzed by flow cytometry. For proliferation, CD4^+^ T cells were stained with CTV and stimulated as described above. CTV expression was analyzed by flow cytometry after 5 days of culture. Results are expressed as the percentages of inhibition of the activation and of the proliferation. **(B)** Comparison of naive (CD44^-^CD62L^+^) and effector/memory (CD44^+^CD62L^-^) CD4^+^ T cell frequencies between a young NZB/W and an old NZB/W mouse by flow cytometry is shown as an example and percentage of CD4^+^ T cell subsets in young (n=13) and old (n=10) NZB/W mice are represented. **(C)** Purified naive CD4^+^ T cells from old BALB/W (blue bars, n=14) and old NZB/W mice (green bars, n=10) were stimulated as in (A) and the percentages of inhibition of the activation and of the proliferation are represented. **(D)** Purified total CD4^+^ T cells (left) or naive CD4^+^ T cells (right) of old BALB/W (blue bars, n=16) and old NZB/W mice (green bars, n=10) were stimulated with anti-CD3/anti-CD28 antibody-coated beads in the presence of the agonist anti-BTLA 6F7 antibody or its isotype control and cultured for 24h. BTLA expression was analyzed by flow cytometry and the BTLA fold enhancement (expressed as a ratio of BTLA MFI following activation/BTLA MFI without stimulation) is represented. **p<0.01; ***p < 0.001; ****p<0.0001, Mann-Whitney test.

If the proportion between naive and effector/memory CD4^+^ T cells did not significantly differ with age in BALB/W mice, there is a strong diminution of naive CD4^+^ T cells (80±2% *vs* 15±3%; p<0.0001) towards an enhancement effector/memory CD4^+^ T cells (20±2% *vs* 85±3%; p<0.0001) in old NZB/W mice compared to young lupus mice (Fig. 2B). We analyzed whether defective BTLA signaling in old lupus mice was associated with this shift of CD4^+^ T cells toward an effector/memory phenotype. When purified naive CD4^+^ T cells from old-diseased NZB/W mice were used in the functional tests, the defective BTLA functionality was not observed anymore (Fig. 2C). Accordingly, BTLA upregulation after activation, which plays an important role to allow this co-inhibitory receptor to further regulate lymphocyte activation, is significantly lower in total CD4^+^ T cells (as we previously demonstrated in lupus patients; 13) but not naive CD4^+^ T cells from old-diseased NZB/W mice compared to control mice of the same age (p<0.01, Fig. 2D). Unfortunately, we were not able to perform the reverse experiment as memory CD4^+^ T cells from old NZB/W mice are refractory to *in vitro* activation (Supplementary Fig. 2). Altogether, our results suggest that defective BTLA functionality in old-diseased NZB/W mice is related to the high frequency of effector/memory CD4^+^ T cells, that may be refractory to BTLA-mediated inhibition.

### Administration of the 6F7 anti-BTLA antibody prevents the development of proteinuria and extends survival

Our phenotypic and functional analyses revealed numerous similarities with data we obtained in lupus patients (13), highlighting the relevance of using this mouse model for evaluating the potential therapeutic effect of targeting the BTLA pathway in lupus. We administered the 6F7 antibody (or its isotype control) into 20-22-week-old NZB/W mice, by the i.p route at 3mg/kg twice a week for 10 weeks and mice were followed for various parameters until they die (Fig. 3A). Typically, dosing for prophylactic treatments in NZB/W begins at 12-14 weeks of age. Therefore, we administered the anti-BTLA 6F7 antibody into NZB/W mice aged 20-22 weeks which already exhibit mild to high levels of autoantibody titers, without any overt pathological manifestations (20). NZB/W mice aged 20-22 weeks can thus be defined as being in the preclinical phase of autoimmunity establishment and not considered as naive animals.

**Figure 3:**
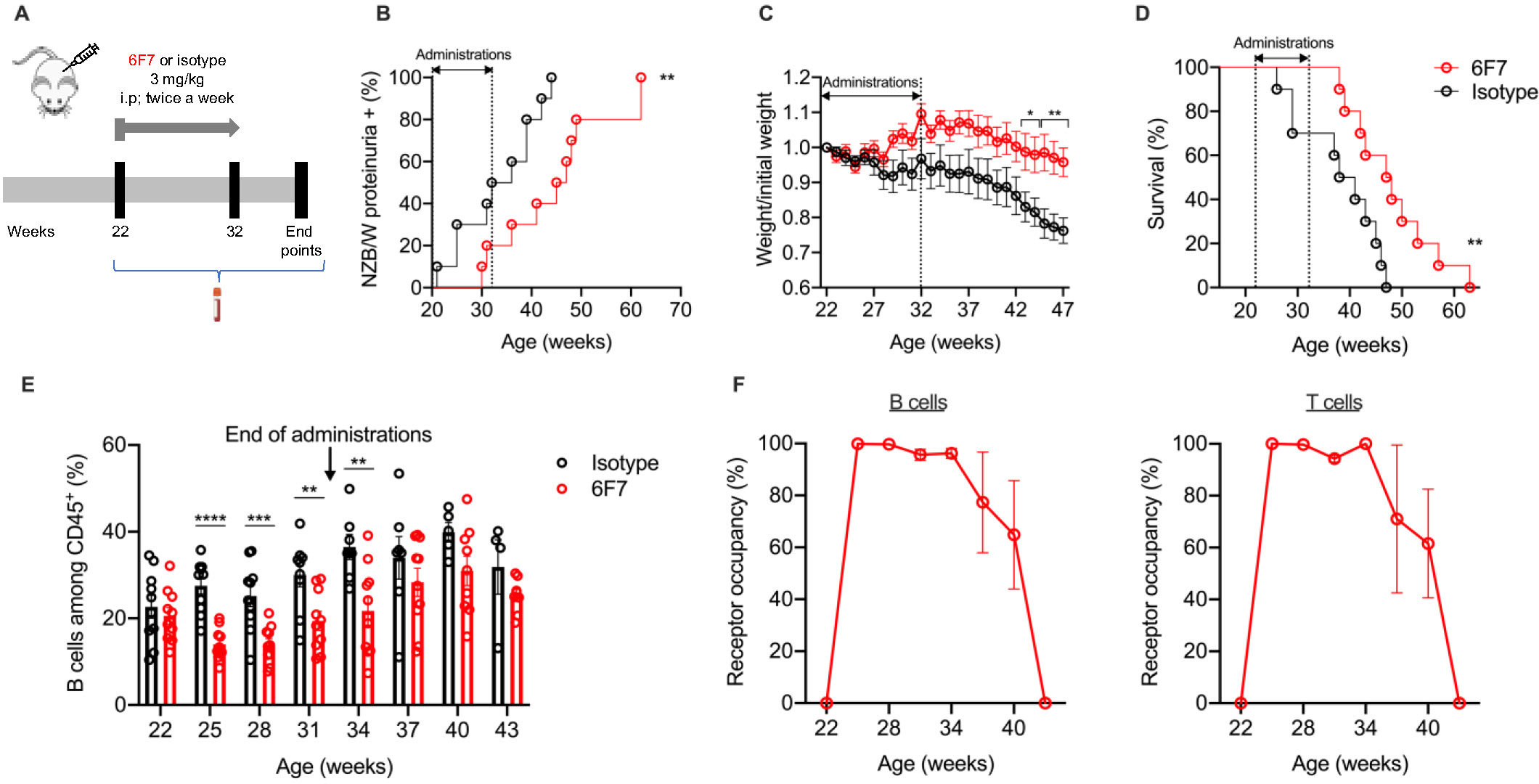
Administration of anti-BTLA 6F7 antibody to NZB/W mice delays the onset of proteinuria and prolongs survival. **(A)** NZB/W mice of 20-22 weeks of age received either i.p. administration of an anti-BTLA antibody (clone 6F7; 3mg/kg/administration, n=10) or an appropriate mouse isotype antibody (IgG1κ; 3mg/kg/administration, n=10) twice a week for 10 weeks. Mice were followed for various parameters and sacrificed at limit points. **(B)** Percentages of NZB/W mice that developed severe proteinuria in the group of anti-BTLA-treated mice (in red) and isotype-treated mice (in black). **(C)** Body weight loss expressed as a ratio compared to the initial body weight in anti-BTLA-treated mice (in red) and isotype-treated mice (in black). **(D)** Percentages of mice that had survived in anti-BTLA treated (red) or isotype control treated mice (black). **(E)** Percentages of peripheral CD19^+^ B cells assessed by flow cytometry in the anti-BTLA-treated group (red) and the isotype-treated group (black). **(F)** Blood B and T cells were stained with a PE-conjugated 6F7 antibody to evaluate receptor occupancy (RO) in 6F7-treated mice during the course of the treatment. RO for each timepoint was calculated as follow: 1-[MFI 6F7/MFI 6F7at day 0]x100. *p<0.05; **p<0.01; ***p<0.001; ****p<0.0001, Log-rank (B, D) or Mann-Whitney test (C, E).

The 6F7 antibody administration into 20-22-week-old NZB/W mice results in delayed proteinuria onset (p<0.01; Fig. 3B). At the age of 44 weeks, all isotype-treated NZB/W mice had developed proteinuria, whereas only 40% of 6F7-treated mice were proteinuria positive. The 6F7 administration also limits the disease-related body weight loss of NZB/W lupus mice (Fig. 3C). More importantly, 6F7-treated mice exhibited an extended survival time (p<0.01; Fig. 3D). The life span of 6F7-treated mice was significantly prolonged and at the age of 47 weeks, all isotype-treated mice were sacrificed according to limit points, whereas 50% of 6F7-treated mice were still alive. The beneficial effect of anti-BTLA 6F7 antibody administrations on lupus development was accompanied by a significant decrease (but not a total depletion) of peripheral CD19^+^ B cells. Indeed, circulating B cell frequencies (but not T cell frequencies; Supplementary Fig. 3) were significantly reduced during the treatment (20.6 ± 1.9% of CD19^+^ B cells before administrations *vs* 14 ± 1.2% after 9 weeks of administrations, p<0.01; Fig. 3E) in anti-BTLA treated mice. Interestingly, the decrease in circulating B cell frequencies perfectly corresponds to BTLA receptor occupancy (RO) which was evidenced by the inability of exogenous labeled 6F7 antibody to bind to CD19^+^ B cells and CD3^+^ T cells from 6F7-treated mice (Fig. 3F). The timing of proteinuria onset and survival enhancement in anti-BTLA treated mice corresponds to the loss of anti-BTLA antibody exposure (10 weeks), suggesting that efficacy requires continuous administration and does not induce long-term protective effect (Fig. 3 B, D).

Unfortunately, we didn’t notice any significant differences concerning levels of circulating anti-dsDNA and anti-chromatin IgG antibodies (data not shown) in comparison with isotype-treated mice.

### Administration of the 6F7 anti-BTLA antibody reduces follicular B cell numbers in the spleen and leucocyte infiltration in the kidneys

In our initial therapeutic protocol, all mice were euthanized when they had reached limit points, making difficult the comparison between isotype and 6F7-treated mice (that even if latter, had developed lupus symptoms) in organs. We initiated a second therapeutic protocol in which mice were euthanized one week after the last administration to evaluate the consequence of BTLA targeting in the spleen and the major targeted organ, i.e. kidneys (Fig. 4A). At the time of sacrifice (i.e at the age of 33 weeks), accordingly to the results obtained in our previous experiment (Fig. 3B), 7 out of 11 mice that had received the isotype displayed proteinuria (64%) whereas only 2 out of 10 (20%) mice treated with the 6F7 antibody were proteinuria positive (Fig. 4B). We also confirmed the significant decrease of B cell frequencies in the blood, except for one mouse (mouse NZB/W1, highlighted with a green frame), suggesting a non-responder status for this mouse (Fig. 4C). The 6F7 administration does not only modify peripheral B cell proportions but also significantly decreases B cell numbers in the blood as evidenced by B cell counts 7 days after 6F7 administration (Supplementary Fig. 4). Splenomegaly, a typical manifestation of lupus progression in the NZB/W mouse model, is also reduced in 6F7-treated mice (total splenocytes numbers were decreased in 6F7-treated lupus mice; Fig. 4D; p=0.07 or 0.04 without the NZB/W 1 mouse). Accordingly, the total numbers of lymphocytes in the spleen seems to be decreased in 6F7-treated mice compared to isotype-treated mice (0.6×10^8^±0.1 *vs*

**Figure 4:**
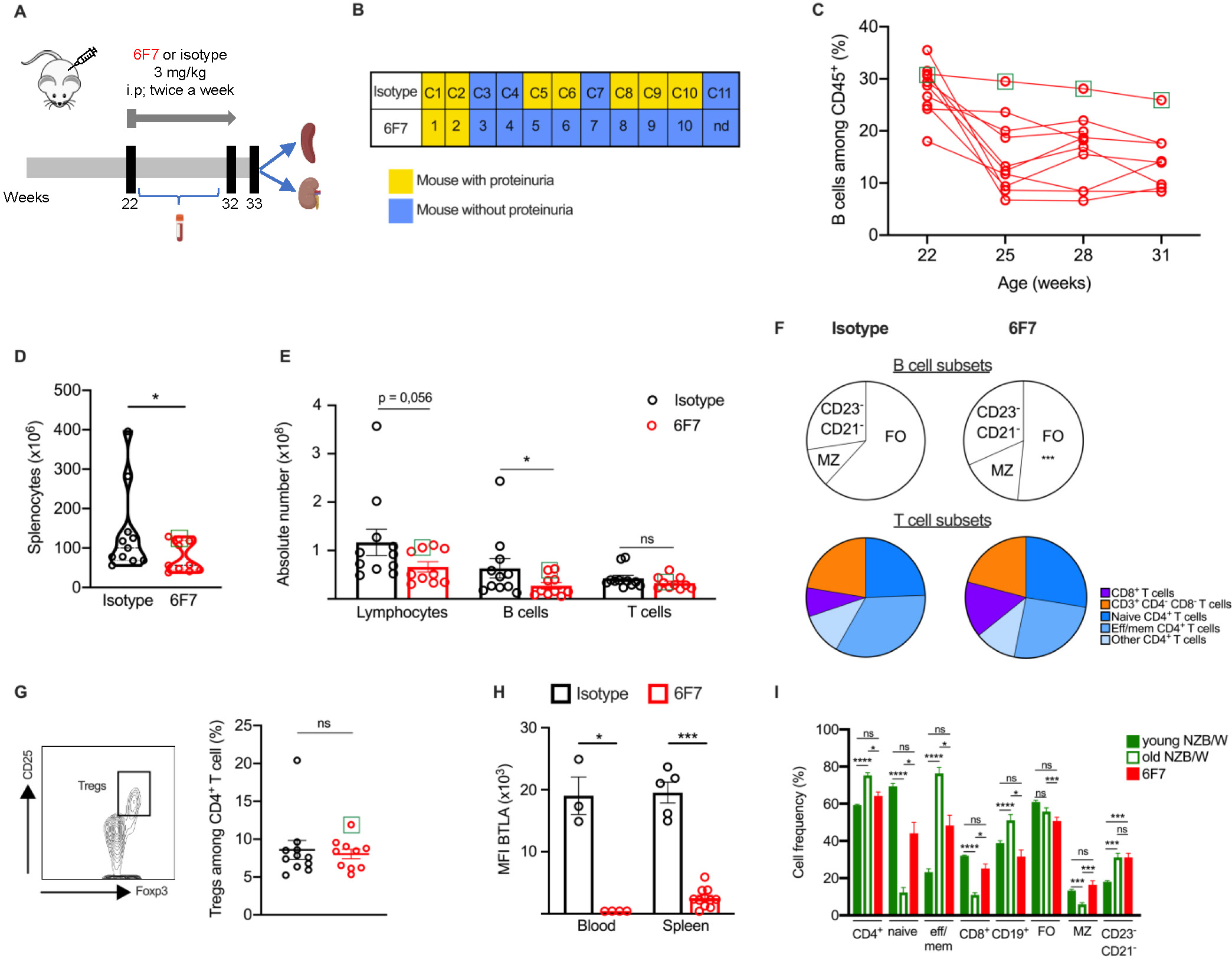
Anti-BTLA 6F7 antibody administration decreases peripheral and spleen FO B cell numbers. **(A)** NZB/W mice of 20-22 weeks of age received either the 6F7 antibody (n=10) or its isotype (n=11) as in Fig. 3 and were sacrificed one week after the end of the treatment. **(B)** Proteinuria status of mice is indicated for each mouse. **(C)** Percentages of peripheral CD19^+^ B cells assessed by flow cytometry in the anti-BTLA-treated group. The NZB/W1 mouse is highlighted with a green frame. **(D)** Comparison of total splenocyte numbers between isotype (in black) and 6F7 (in red) treated mice **(E-F)** Absolute numbers of total lymphocytes, B and T cells (**E**), as well as B and T cell subset frequencies (**F**) in the spleen were determined in mice that were administered (for 10 weeks) with the 6F7 antibody (in red) or its isotype control (in black). **(G)** Frequency of Tregs (CD25^+^Foxp3^hi^) among CD4^+^ T cells in the spleen of isotype (in black) and 6F7-treated mice (in red). **(H)** Mice were administered with 2×3mg/kg of 6F7 antibody (in red) or isotype (in black) at day 1 and day 4, and were sacrificed on day 7 to collect blood and spleens. RO was determined thanks to PE conjugated anti-BTLA (6F7) staining. (**I**) Comparison of lymphocyte frequencies in the spleen of non-diseased NZB/W mice (12 wk-old), diseased NZB/W mice (proteinuria positive, >35-week-old) and 6F7-treated NZB/W mice. Each dot represents one mouse. *<0.05; ***p<0.001; ****p<0.0001, Mann-Whitney (D-H) or Kruskal-Wallis with post hoc analysis (Dunn’s test) (I). The non-responder NZB/W1 mouse is highlighted with a green frame and was excluded of the analysis.

1.2×10^8^±0.3; p=0.09 or p=0.056 when the NZB/W1 mouse was excluded from the analysis; Fig. 4E). The reduced lymphocyte numbers concerns B cells (p=0.072 and 0.031 without the NZB/W1 mouse) and particularly FO B cells (those expressing very highest levels of BTLA; p<0.001) whereas T cell numbers remain unchanged (Fig. 4E, F). Contrary to what have been shown in the NOD mouse model (21), we did not notice any Treg expansion in the spleens of NZB/W lupus mice following 6F7 administrations (Fig. 4G). The discreet effect of 6F7 administrations on splenocytes (compared to the high reduction of peripheral B cell frequencies) is not due to the inefficiency of the 6F7 antibody to penetrate the spleen as BTLA detection with exogenous labeled 6F7 antibody is decreased to the same degree than in the blood (Fig. 4H). Finally, we observed that immune cell proportions in the spleens of 6F7-treated mice, are more closely related to those of young, non-diseased NZB/W mice (12-week-old) than those of old-diseased animals (>35-week-old and proteinuria positive), suggesting that the 6F7 administration was able to restore immune cell alterations occurring during the development of the disease (Fig. 4I).

As proteinuria is a marker of renal lesions, we performed histological analyses on kidneys from 6F7-treated mice. Accordingly with the results obtained in collected urine samples, anti-BTLA treated mice displayed limited kidney damages with a reduction in glomerular size and cellularity, mesangial expansion and deposits, and glomerular damages, all taken into consideration in the histopathological score (p<0.05; Fig. 5A). Kidneys of anti-BTLA-treated mice were also less infiltrated by immune cells. Indeed, the kidney infiltrate area is twice lower in 6F7-treated mice compared to isotype-treated mice (no mouse displaying an area of infiltrate higher than 2 compared to 6 mice out of 11 in the control group; Fig. 5B), even if the comparison is not statistically significant, probably because of the great heterogeneity observed in the isotype group (p=0.13 and p=0.07 without the NZB/W1 mouse). Indeed, as depicted in Fig. 4B, 4 mice of the isotype-treated group (mice C3, C4, C7 and C11) did not exhibit any clinical manifestations at the time of their sacrifice and accordingly are those for which we did not observe any CD45^+^ cell infiltration in the kidney. FACS-analysis confirmed the decreased frequencies of CD45^+^ cells in the kidneys of anti-BTLA treated mice (Fig. 5C p<0.05) with reduced infiltration of effector/memory CD4^+^ T cells (Fig. 5D, ns but p=0.03 if the NZB/W1 mouse is not included in the analysis) and a tendency to decrease in plasma cell frequencies (p=0.09, p=0.06 without the NZBW1; Fig. 5E).

**Figure 5:**
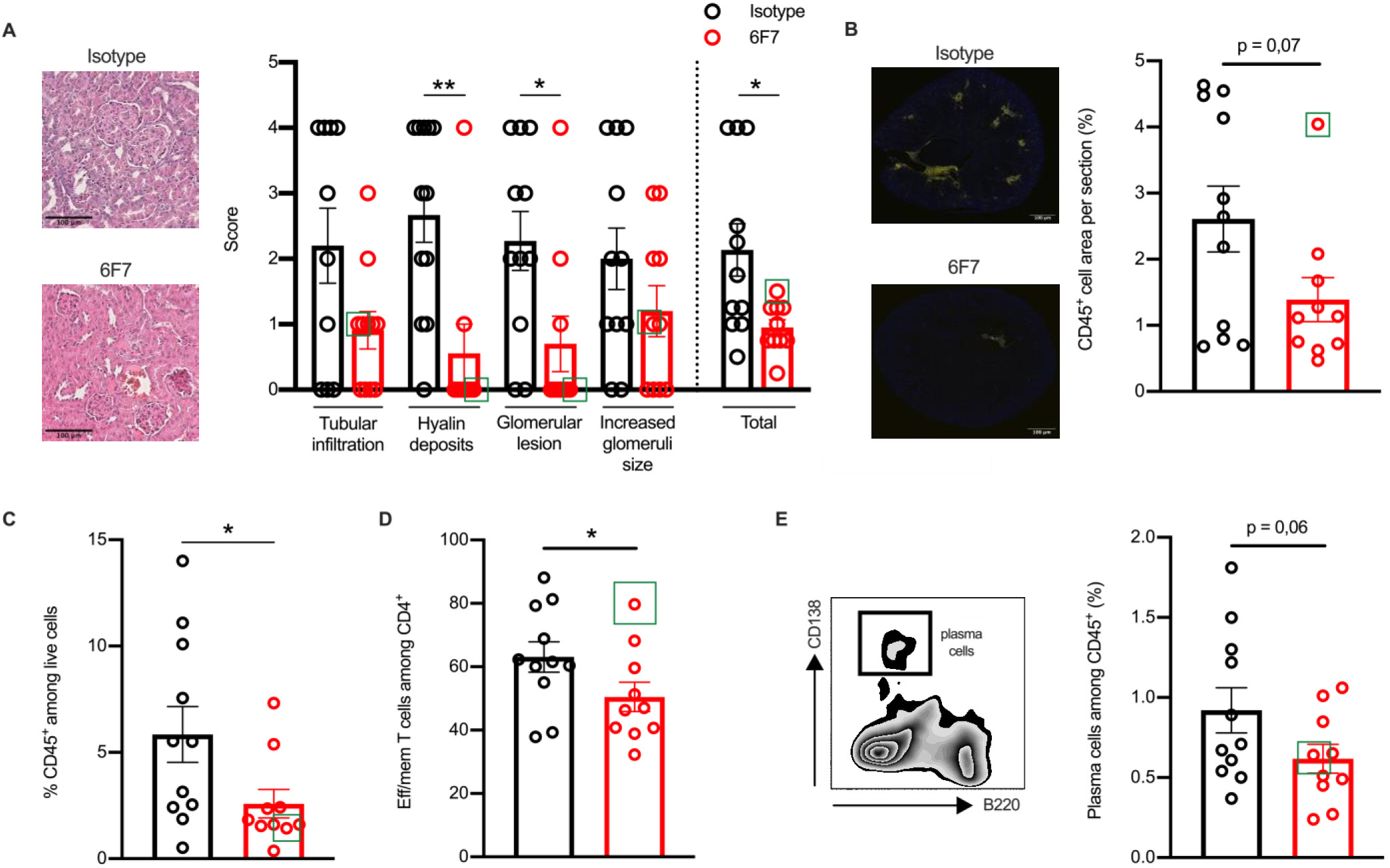
Anti-BTLA 6F7 antibody administration reduces kidney damages. 20-22 NZ/W mice received either the 6F7 antibody (n=10) or its isotype (n=11) as in Fig. 3 and were sacrificed one week after the end of the treatment. **(A)** Kidney sections from a representative isotype-treated NZB/W or an anti-BTLA-treated NZB/W mice (left) and comparison of pathological changes (right) assessed by evaluating glomerular activity and tubulointerstitial activity between anti-BTLA (red) and isotype (black) treated mice. **(B)** Kidney sections from anti-BTLA 6F7 or isotype-treated mice were stained with anti-CD45 antibodies (yellow). Entire sections were reconstituted using Metamorph software and areas corresponding to tubulointerstitial infiltrates were quantified with ImageJ software. Results are expressed as the ratio between the surface of infiltrates and the total surface of the kidney as a percentage. Scale bars: 100µm; (original magnification x100). Representative images of one section from one mouse of each group (6F7 or isotype-treated mice) is shown. (**C-E**) FACS-analysis of leukocytes (**C**), effector /memory CD4^+^ T cells (**D**) and plasma cells (**E**) frequencies in 6F7 mice (in red) or isotype-treated mice (in black). Each dot represents one individual. *p<0.05; **p<0.01, Mann-Whitney test. The non-responder NZB/W1 mouse is highlighted with a green frame and was excluded of the analysis.

### Administration of 6F7 down-modulates BTLA cell surface expression and induces B cell hyporesponsiveness

Antibody targeting surface molecules can lead to different outcomes such as receptor down modulation, depletion, blocking of receptor/ligand interaction or agonist activities. To analyze whether the 6F7 antibody administration modifies BTLA expression, we used another anti-BTLA antibody (clone 8F4), which recognizes another epitope than the 6F7 clone as confirmed by the double staining (Fig. 6A, top panel) and which is still able to bind BTLA following 6F7 exposure (Fig. 6A, bottom panel). BTLA expression is drastically decreased on peripheral B cells (Fig. 6B) whereas it remains unchanged on peripheral T cells (Fig. 6C). In the spleen, we observed a reduced but still detectable BTLA expression on B cells (particularly FO B cells; Fig. 6D) and only a slight diminution on T cells (Fig. 6E; only significantly reduced on CD4^+^ T cells) from 6F7-treated mice. Surprisingly, we observed a decrease of BTLA expression on peripheral B cells from the NZB/W1 mouse (Fig. 6B) but not by its spleen B cells (Fig. 6D). BTLA downmodulation is not due to its internalization following 6F7 binding, as we did not observe any decrease in BTLA detection (with the 8F4 clone) when splenocytes were incubated at 37°C with saturating amounts of 6F7 at various time compared to incubation at 4°C (Fig. 6F), although the 6F7 antibody was efficiently bound to BTLA as revealed by the fluorescent-6F7 staining. Moreover, the decreased of BTLA expression in 6F7-treated mice compared to isotype-treated mice is similar when we compared surface and intracellular staining, indicating BTLA is not more sequestered into the cells of mice that had received the 6F7 antibody (Fig. 6G).

**Figure 6:**
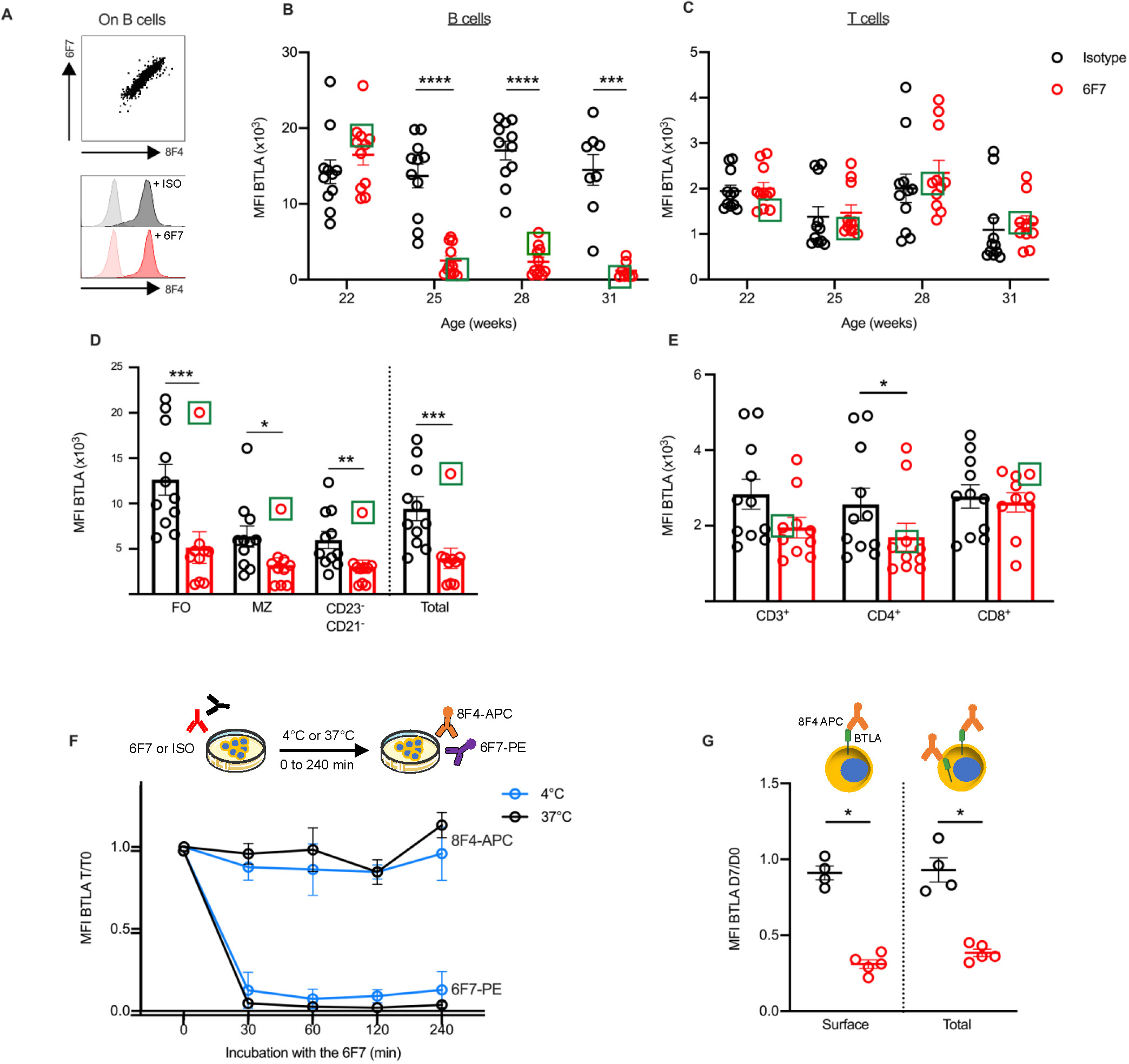
Administration of anti-BTLA 6F7 induces down modulation of BTLA expression. **(A)** The 8F4 anti-BTLA antibody can be used to detect BTLA expression following 6F7 exposure. Top: representative image of spleen cells co**-**stained with APC conjugated anti-BTLA (8F4) and PE conjugated anti-BTLA (6F7). Bottom: spleen cells were incubated with saturating amounts of unlabeled anti-BTLA 6F7 (10µg/ml; in red) or isotype control (in black), and then stained with anti-CD19 and anti-BTLA (clone 8F4) antibodies. FMO for each condition are indicated. **(B-C)** BTLA expression (detected by the 8F4 antibody) on peripheral B cells (gated on CD45^+^CD19^+^ cells) and peripheral T cells (gated on CD45^+^CD3^+^ cells) of isotype (in black) or 6F7-treated mice (in red). **(D-E)** BTLA expression (detected by the 8F4 antibody) on B cells (gated on CD45^+^CD19^+^ cells; D) and T cells (gated on CD45^+^CD3^+^ cells; E) isolated from the spleen of isotype (in black) or 6F7-treated mice (in red) sacrificed one week after the last administration (33-week-old). **(F)** Spleen cells from BALB/W mice (n=3-4) were incubated with saturating amounts of 6F7 antibody (10µg/ml) for 30, 60, 120 and 240 minutes at 4°C or 37°C and then stained with the APC conjugated 8F4 or the PE conjugated 6F7 antibodies. Results are expressed as ratio of MFI (T/ T0). **(G)** Mice were administered with 2×3mg/kg of 6F7 antibody (n=5 in red) or isotype (n=4 in black) at day 1 and day 4, and the blood was collected on day 7. Membrane (surface BTLA) and intracellular (total BTLA) staining with the 8F4 antibody were performed and BTLA expression is shown. Results are expressed as ratio of MFI between day 7 and day 0. Each dot represents one individual. *p<0.05; **p<0.01; ***p<0.001; ****p<0.0001 Mann-Whitney test.

To address whether the 6F7 could deplete BTLA^+^ cells *in vivo*, we administered mice with a very high dose of the 6F7 antibody (500µg; 15mg/kg) and analyzed lymphocyte numbers in the spleen 5 days later. To decipher with ADCP or ADCC involvement, some mice were depleted in macrophages or NK cells prior 6F7 administrations. No significant decrease in lymphocyte numbers (CD19^+^ B cells or CD3^+^ T cells) in the spleen 5 days after the i.p. injection of 6F7 was observed (Fig. 7A; p=0.4 for B cells and p>1 for T cells), which agreed with the lack of depleting activity reported for the majority of mouse antibodies of the IgG1 isotype. Moreover, spleen cells incubated in the presence of the 6F7 antibody did not die from apoptosis (Fig. 7B). We also explored whether 6F7 could act as a blocking antibody and found that the binding of a HVEM fusion protein to BTLA is preserved when spleen cells were incubated with saturating amounts of the 6F7 antibody (Fig. 7C). Finally, we wondered whether the 6F7 administration influences B cell capacity to be activated. We analyzed CD69 expression on B cells isolated from mice that had received either the 6F7 antibody or the isotype control, and evidenced that B cells from 6F7-treated mice are less responsive to BCR activation (Fig. 7D), indicating an agonist activity of the 6F7 antibody *in vivo*. Altogether, our results show that the 6F7 antibody, is not a depleting antibody, induces a decrease of BTLA expression particularly on cells expressing high levels of BTLA, does not block the HVEM binding to BTLA and reduces B cell activation.

**Figure 7:**
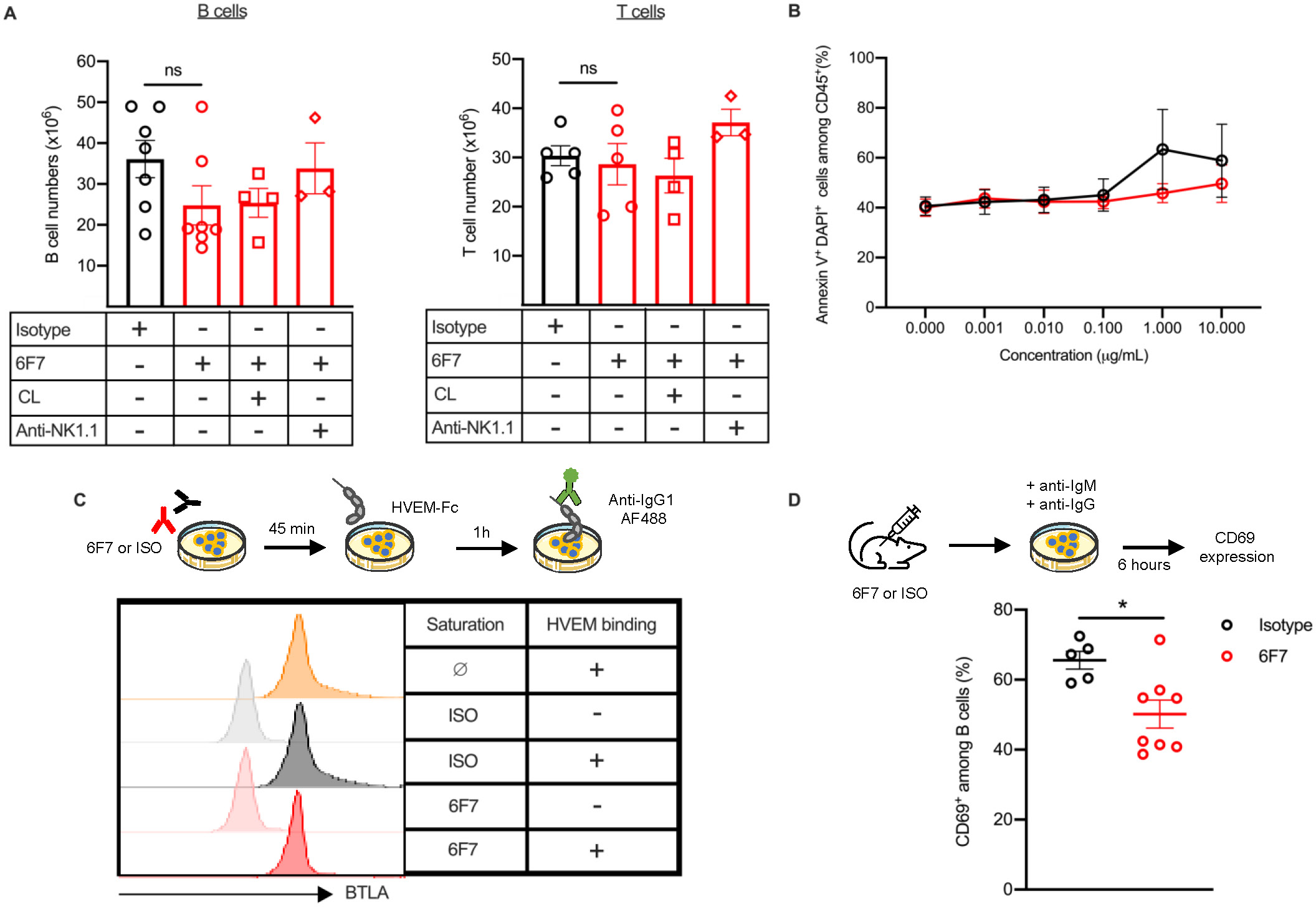
Activation of BTLA leads to B cell hyporesponsiveness. **(A)** Mice were administered with 500 µg of 6F7 antibody (15mg/kg; in red, n=7) or isotype (in black, n=7) at day 0 and were sacrificed on day 6 to collect spleens and determined cell numbers. Some mice were firstly depleted in NK cells thanks to i.p. administrations of an anti-NK1.1 depleting antibody (clone PK136; 200µg on days −2, 0 and 3, n=3) or in macrophages thanks to i.v. administration of clodronate-loaded liposomes (200µl on day −1, n=4). **(B)** Spleen cells were incubated with increasing concentrations of 6F7 or isotype antibodies for 20-24h. Cell death was measured with DAPI and Annexin-V staining on CD45^+^ cells. As control, apoptosis was also analyzed in the presence of H_2_O_2_ (that gives >90% of cell death). **(C)** Spleen cells were firstly incubated with saturating amounts of unlabeled 6F7 (10µg/ml) or isotype antibodies for one hour at 4°C. After washing, cells were incubated with HVEM-Fc mouse fusion protein (20 µg/mL) for 1h at 37°C and binding of HVEM-Fc to BTLA was revealed with an AF488-conjugated anti-IgG1secondary antibody. One representative experiment of 4 with similar results is depicted. **(D)** Spleen cells isolated from mice treated as in (**A**), were stimulated with anti-IgM and anti-IgG antibodies for 6h and the expression of CD69 was evaluated by cytometry. Each dot represents one mouse. *p<0.05, ***p<0.001, Mann-Whitney.

## DISCUSSION

Upon its discovery, research into the involvement of the BTLA pathway in autoimmune disorders has swiftly emerged. However, despite mounting evidence underscoring BTLA’s role in maintaining tolerance, its implication in lupus pathogenesis remains unclear. While recent studies have investigated BTLA expression and function in human patients’ cells (13–14, 22), only one study in a mouse lupus model has shown that BTLA deficiency exacerbates lupus disease progression (12). In the present study, we conducted a comprehensive analysis of BTLA expression and function in the classical NZB/W lupus-mouse model. Our findings reveal no significant alteration in BTLA expression compared to control mice, except an enhancement on CD21^-^CD23^-^ and MZ B cells particularly in aged animals. This higher BTLA expression could reflect an activated state of these cells, as activated cells typically exhibit increased BTLA levels. Moreover, CD21^-^CD23^-^ express several markers characteristics of memory activated cells including Fas, CD80, CD86 or MHC class II molecules (23).

Additionally, we found that akin to human lupus T cells, the BTLA signaling pathway is impaired in CD4^+^ T cells from diseased-NZB/W lupus mice. Although we couldn’t assess BTLA functionality in purified effector/memory CD4^+^ T cells from diseased NZB/W mice due to their poor proliferative response compared to naive T cells upon antigenic stimulation (24 and this work), our findings unequivocally show that the compromised BTLA functionality observed in total CD4^+^ T cells is not attributed to naive CD4^+^ T cells. In lupus patients, we evidenced that the impaired BTLA functionality is due to defective recruitment of BTLA to preexisting TCR clusters. Given that TCR clusters have been observed even without stimulation in diseased NZB/W mice (25), we hypothesize that altered BTLA functionality in these NZB/W mice is linked to the exclusion of BTLA within these pre-clustered TCR in effector/memory T cells from diseased animals.

Furthermore, we investigated the impact of administering an anti-BTLA antibody on disease features in NZB/W mice, revealing compelling evidence that targeting this pathway may have disease-modifying activity. Remarkably, *in vivo* treatment with the anti-BTLA 6F7 antibody significantly decreased mortality rate among NZB/W mice and mitigated renal infiltration by lymphocytes, consequently delaying the onset of nephritis. Relapse occurred rapidly after the loss of BTLA receptor occupancy, suggesting that the antibody’s beneficial effect probably does not involve the induction of a tolerance mechanism but instead requires ongoing stimulation of BTLA pathway. Consistent with this, we did not notice any modification in Treg frequency in 6F7-treated animals. Contrary to what we could expect, administration of the 6F7 did not result in reduced levels of anti-dsDNA or anti-nucleosome auto-antibodies. However, previous findings in the MRL/lpr lupus model demonstrated that B cells alone (rather than autoantibodies) are sufficient for nephritis and cell infiltration (26). Accordingly, several studies have shown therapeutic strategies to be beneficial in lupus mice without concomitant decreases in anti-DNA antibody levels (27–28). Conversely, in certain cases, diminution of autoantibodies did not translate to improvements in pathology (29). Similarly, in the Ldlr^-/-^ model, administration of the agonist anti-BTLA 3C10 antibody (7) attenuates atherosclerosis, without affecting total or antigen-specific antibodies levels, aligning with data showing that allogenic humoral response remains unaffected following antibody-mediated blockade of the HVEM/BTLA pathway (30).

Some of autoantibodies are produced by long-lived plasma cells (LLPCs), which are established very early in the spleen of NZB/W lupus mice as demonstrated by previous research (31). LLPCs can be detected as early as 6 weeks of age and their numbers continue to increase until they reach a plateau at 10 weeks of age. However, LLPCs only begin to appear in the kidneys at 16 weeks of age, and their numbers constantly increase throughout the progression of the disease. We found that while frequency of CD138^+^ plasma cells decreases in the kidneys of 6F7-treated mice, it remains unchanged in the spleens (data not shown). This suggest that although 6F7 treatment does not significantly impact the well-established LLPC in the spleen, it appears to limit their accumulation in kidneys.

Published data on the mode of action (MOA) of anti-BTLA antibodies *in vivo* vary depending on the clone that was used: the 6A6 antibody is described as a blocking but non-depleting antibody (8), while clones 3C10, BYK-1, 4G12b are classified as agonist antibodies (6, 32–33). Contradictory data have been reported in the literature regarding the 6F7 antibody: Truong et al documented that the 6F7 antibody depleted B and T cells in the NOD diabetic model (19) whereas it exhibited *in vivo* agonist activities in an induced-colitis model and in tuberculosis (34–35). Depleting antibodies can induce cell death through various mechanisms, including complement-dependent cytotoxicity (CDC), direct induction of apoptosis and antibody-dependent cellular cytotoxicity or phagocytosis (ADCC/ADCP). The 6F7 antibody is an IgG1 isotype, which is unable to bind C1q and therefore does not efficiently activate the classical complement pathway. Furthermore, we also evidenced that the 6F7 antibody’s inhibition of B cells does not involve induction of apoptosis. Lastly, it’s worth noting that mouse IgG1 does not bind the mouse FcRI and FcRIV which are the primary Fc receptors involved in ADCC or ADCP mediated by NK and myeloid cells (36). This limits the likelihood that the 6F7 antibody might have induced depletion trough ADCC mechanism, which was confirmed by our results. B cell depletion from the circulation could potentially involve mechanisms beyond FcgR-mediated phagocytosis. It might also reflect B cell migration into tissues rather than B cell destruction. Therefore, the drastic observed reduction of peripheral lymphocytes in lupus mice treated with the anti-BTLA 6F7 antibody might thus be due to non-depletive processes.

Our results clearly demonstrate that administering the 6F7 antibody results in diminished BTLA expression in peripheral and spleen lymphocytes. The 6F7 MOA thus resembles to the one of the anti-BTLA clone PJ196, which did not deplete BTLA expressing cells but caused its down regulation, promoting islet allograft acceptance (37). The BTLA downmodulation precisely correlates with the decline in cell frequency: among the cell populations, FO B cells exhibit the most substantial reduction in BTLA, and notably, they are the sole cell population wherein a significant decrease in numbers was observed in the spleen. The marked reduction in BTLA levels, coupled with the decrease in the number of blood B cells, primarily comprising FO B cells circulating though the blood, further corroborate that FO B cells represent the primary target of the 6F7 antibody. Accordingly, it was suggested that BTLA expression was necessary for the survival of FO B cells (29). In line with our findings, the administration of the agonist 3C10 anti-BTLA antibody also selectively reduced FO B cells in the Ldlr^-/-^ mouse model of atherosclerosis (7). Interestingly, and not surprisingly given that BTLA is a negative regulator of Syk (38), the oral administration of a selective Syk inhibitor attenuates lupus nephritis and also reduces FO B cells without affecting the proportion of MZ B cells in the NZB/W lupus model (20).

The scenario involving the NZB/W1 mouse presents an intriguing puzzle. Although the 6F7 staining wasn’t assessed in this mouse, the observed reduction in BTLA expression on peripheral B cells observed with the 8F4 clone (as depicted in Fig 6B) indicates efficient binding of the 6F7 antibody to BTLA. Despite this effective binding on peripheral B cells, there’s no reduction in B cell frequencies in the blood, suggesting that the mouse doesn’t react to the anti-BTLA treatment, as also evidenced by the absence of effect in all clinical parameters. Moreover, BTLA expression remains unchanged in spleen B cells of this mouse (contrary to other mice), hinting that the treatment’s efficacy may hinge on the antibody reaching the spleen. Altogether, our data provide evidence that BTLA downmodulation is part of the MOA of the 6F7 antibody.

Alternatively, the 6F7 antibody may affect survival of multiple cell type by modulating BTLA activity in a non-competitive manner with its ligand HVEM. Indeed, BTLA also serves as a ligand to deliver pro-survival cosignals (30). Consequently, as BTLA expression diminishes following 6F7 exposure, fewer positive signals are engaged to HVEM-expressing cells. Additionally, the binding of HVEM is preserved in the presence of the 6F7 antibody. It does not block the engagement of HVEM, thus allowing the endogenous inhibitory pathway to remain active. Besides reducing BTLA expression and FO B cells, we found that 6F7 administration significantly reduces B cell responsiveness to BCR stimulation. This indicates that, in addition to its agonist activity on CD4^+^ T cells, the 6F7 antibody acts *in vivo* through direct agonism of the BTLA pathway on B cells.

Current treatments for lupus do not adequately control disease activity and tissue damages and are associated with significant side effects. Therefore, the identification of new therapeutic targets is a major unmet clinical need. We suggest that utilizing of a non-depleting anti-BTLA antibody may provide an improved safety profile by avoiding issues related to immune suppression. Unlike anti-CD20 antibodies, which depletes all B cell subsets, the 6F7 antibody specifically reduces but not ablates FO B cells and attenuates without entirely halting B cell activation. This mechanism appears to address immune abnormalities that contribute to lupus symptoms. Therefore, activating the BTLA emerges as promising avenue for modulating lupus disease. Progressing in this direction, Lilly recently completed the recruitment of SLE patients for a phase 2 clinical trial evaluating the safety and efficacy of LY3361237, an agonist anti-human BTLA antibody (22B3 antibody, 39).

In conclusion, our study highlights the pivotal role of BTLA signaling in lupus pathogenesis. To date, there are no report exploring the potential of targeting this pathway in murine lupus. We demonstrated that administering the 6F7 antibody to pre-diseased NZB/W mice alleviates the lupus disease development. The MOA of the non-depleting 6F7 antibody are multifaceted and include *i*) the downmodulation of BTLA expression, likely contributing to the decrease of FO B cell numbers and the reduction of positive signals to HVEM-expressing cells, ii) the preservation of HVEM binding allowing endogenous inhibitory pathway to remain active, iii) and the direct agonist activity.

## ACKNOWLEDGEMENTS

This work was supported by the French Centre National de la Recherche Scientifique (CNRS), the French Society of Rheumatology (grant to FM) and the French “Ministère de l’Enseignement et de la Recherche” (fellowship to LA and LG). We thank Delphine Lamon and Fabien Lhericel (CNRS UP3572) for helping with mouse experiments. We thank Blandine Maître for her generous gift of clodronate-loaded liposomes.

## AUTHOR CONTRIBUTIONS

FM designed the study. LG, LA and MS performed the experiments and analyzed the data. FM wrote the manuscript. JDF assisted with microscopy and reviewed the article. HD and LA participated to discussions and reviewed the article.

## CONFLICT OF INTEREST

The author declare that they have no competing interests.

## Notes

### Competing Interest Statement

The authors have declared no competing interest.

