## supplementary figures for "Targeting B and T lymphocyte attenuator regulates lupus disease development in NZB/W mice"

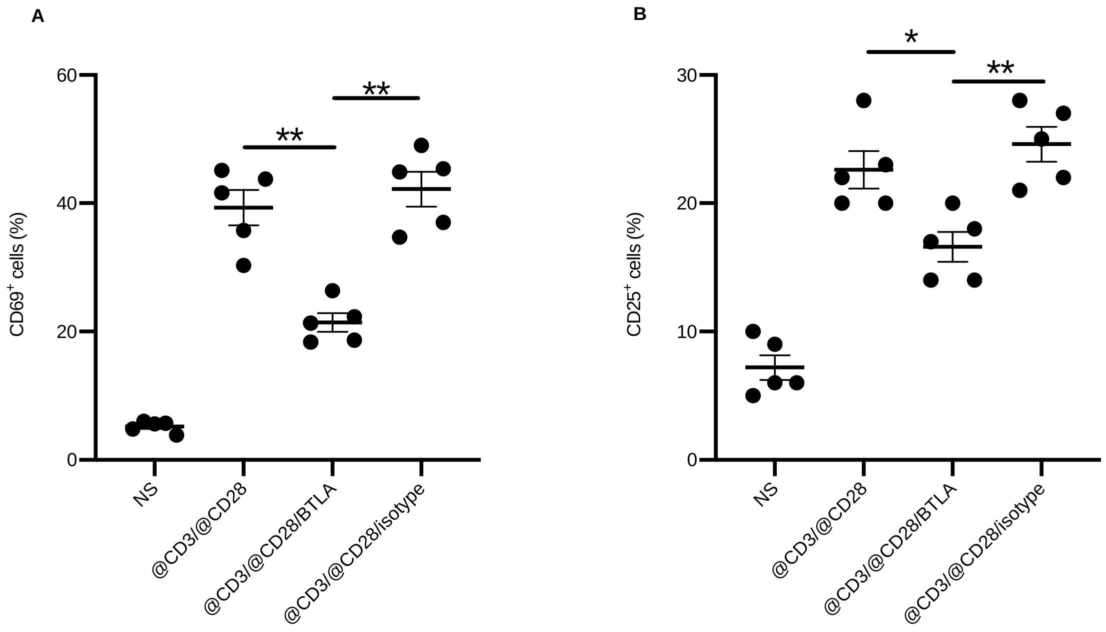


**Supplementary figure 1:** The 6F7 anti-BTLA antibody has an agonist activity in mice carrying the BTLA.1 allele. Percentage of CD69$^{+}$ **(A)** or CD25$^{+}$ cells **(B)** among CD4$^{+}$ T cells from BALB/c mice (n=5). Purified CD4$^{+}$ T cells were stimulated (anti-CD3/anti-CD28 antibody-coated beads) or not (NS) for 24h or 48h in the presence of the agonist anti-BTLA 6F7 antibody or its isotype control (IgG1). CD69 (24h; **A**) and CD25 (48h; **B**) expression were measured by flow cytometry and the percentages of positive cells are shown. *p<0.05; **p<0.01, Mann-Witney test.

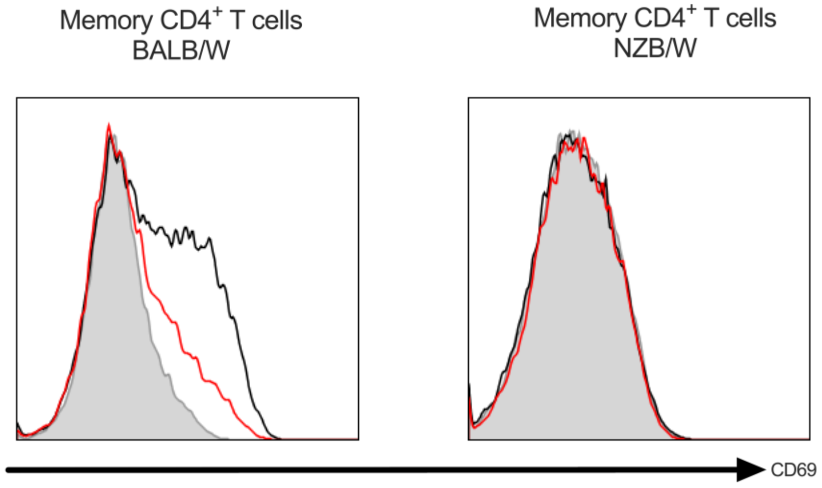


**Supplementary figure 2**: CD4$^+$ T cells from old NZB/W mice are refractory to *in vitro* activation. Purified memory CD4$^+$ T cells from old BALB/W and NZB/W mice were stimulated with anti-CD3/anti-CD28 antibody-coated beads and the expression of CD69 at 24h was analyzed by flow cytometry.

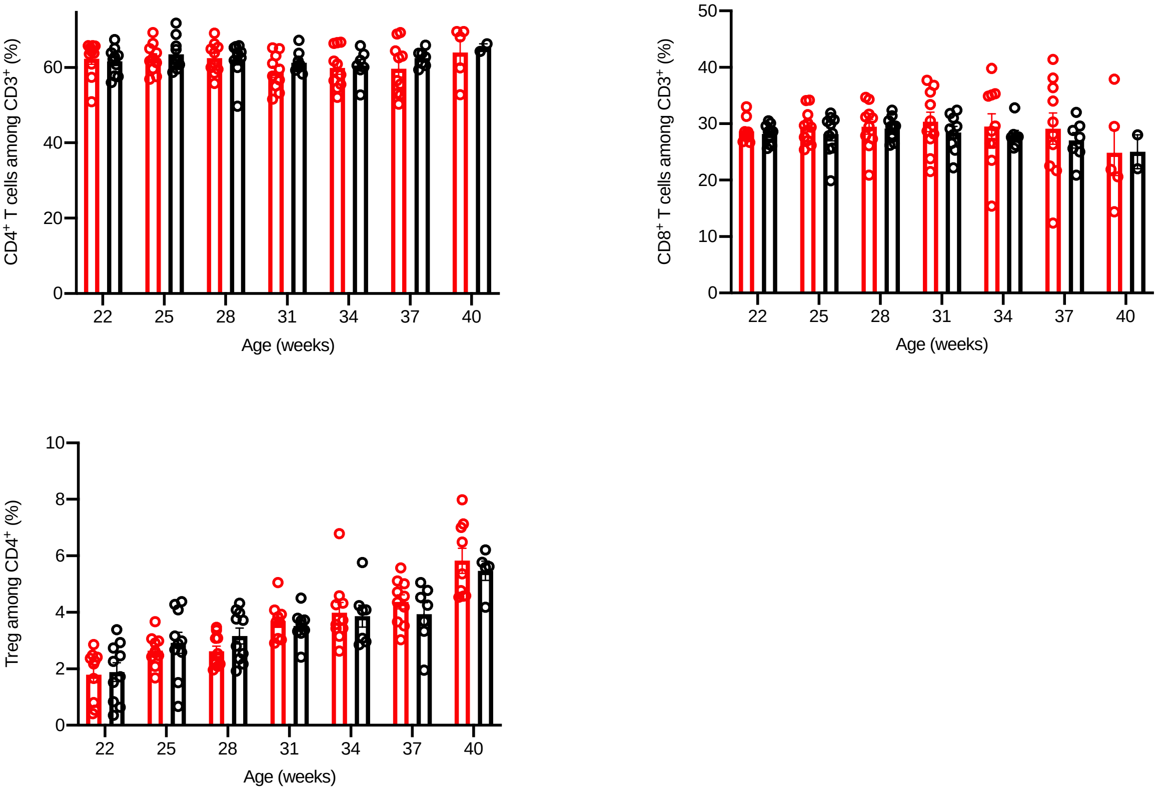


**Supplementary figure 3:** Anti-BTLA 6F7 antibody administrations do not modify peripheral T cell frequencies. Percentages of peripheral $CD4^+$ and $CD8^+$ T cells among $CD3^+$ T cells and of Tregs ($CD25^+FoxP3^{hi}$) among $CD4^+$ T cells were assessed by flow cytometry in the anti-BTLA-treated group (red) and the isotype-treated group (black) every 3 weeks.

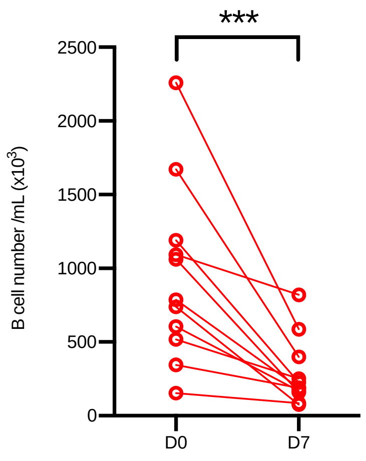


**Supplementary figure 4:** Anti-BTLA 6F7 antibody administration reduces peripheral B cell numbers. Comparison of B cell counts (day 0 and day 7) in mice (n=9) that had received 2 administrations (3mg/kg/administration on day 1 and day 4) of the anti-BTLA 6F7 antibody**.** ***p<0.001, Wilcoxon test.
